# Automated Segmentation of the Hypothalamus: Associations of Age and Socioeconomic Status with Subunit Volumes in Cognitively Healthy Adults

**DOI:** 10.64898/2026.10.31.763278

**Authors:** Barış Akar, Ümit Atasever

**Author notes:** Corresponding author: Barış Akar. Ümit Atasever.

## Abstract

**Background:** The hypothalamus regulates sleep–wake cycles, energy balance, and autonomic and neuroendocrine function, yet its small size and poorly contrasted boundaries have long hindered in vivo volumetry. Deep-learning segmentation now allows the hypothalamus to be divided into functionally meaningful subunits. We examined whether hypothalamic subunit volumes are associated with age and socioeconomic status (SES) in cognitively healthy adults.

**Methods:** From the OASIS-1 cross-sectional dataset (n = 416, 18–96 years), 133 participants with a Clinical Dementia Rating of 0 and complete segmentation output were analysed (33–94 years). A FreeSurfer-based convolutional neural network, run on a cloud virtual machine, segmented five subunits per hemisphere. Volumes were normalised to estimated total intracranial volume (eTIV) and related to age and SES (Hollingshead index) using Pearson correlations with Benjamini–Hochberg false discovery rate (FDR) correction; partial correlations adjusted for sex (age) and for age and sex (SES) served as sensitivity analyses.

**Results:** Older age was associated with lower normalised volume of the right posterior subunit (*r* = −.456, *q* < .001) and the left posterior subunit (*r* = −.274, *q* = .007); both associations persisted after adjustment for sex. Lower SES showed a trend towards larger left anterior-inferior volume (*r* = .242, uncorrected *p* = .005) that did not survive FDR correction (*q* = .053).

**Conclusions:** Age-related volumetric differences in the healthy hypothalamus appear to be concentrated in its posterior portion, a region housing arousal-promoting circuitry. Automated subunit segmentation offers a scalable tool for studying the hypothalamus in ageing and in relation to environmental exposures; the cross-sectional associations reported here should not be interpreted as causal or as within-person change.

## 1. Introduction

The hypothalamus is a small diencephalic structure, occupying well under 1% of brain volume, that orchestrates a remarkably broad set of functions, including the sleep–wake cycle, feeding and energy expenditure, thermoregulation, stress responses through the hypothalamic–pituitary–adrenal (HPA) axis, and innate social behaviours.^1–3^ Its clinical relevance is illustrated by narcolepsy, in which the loss of a small population of hypocretin/orexin neurons in the lateral hypothalamus is sufficient to destabilise sleep–wake regulation.^4^ The hypothalamus has also been proposed as a regulator of systemic ageing: in mice, hypothalamic inflammatory signalling through IKK-β/NF-κB accelerates, and its inhibition delays, age-related decline.^5^

Despite this importance, the human hypothalamus has been comparatively neglected in structural neuroimaging. Its boundaries with neighbouring structures are poorly defined on standard T1-weighted images, manual delineation is time-consuming, and inter-rater variability limits reproducibility and scalability. Billot and colleagues addressed this problem with a convolutional neural network that segments the hypothalamus and five associated subunits (anterior-superior, anterior-inferior, superior tubular, inferior tubular and posterior) per hemisphere, and that is distributed within FreeSurfer.^6,7^ Related deep-learning tools now enable robust segmentation in large and heterogeneous datasets.^8,9^ The subunit approach has since been applied to psychiatric illness, sleep apnoea, childhood conduct problems and even neonates,^10–13^ and large population studies have reported region-specific associations of hypothalamic volume with age, sex and body mass index.^14,15^

Two questions remain incompletely answered. First, it is unclear which hypothalamic subunits differ most with age among cognitively healthy adults; this matters because the subunits contain functionally distinct nuclei—for example, the suprachiasmatic nucleus in the anterior-inferior subunit and the mammillary bodies and arousal-related lateral and tuberomammillary regions in the posterior subunit.^6,16,17^ Second, socioeconomic disadvantage is associated with chronic stress exposure and altered brain structure,^18–20^ yet its relationship with hypothalamic subunit volume has rarely been examined.

Accordingly, we applied automated hypothalamic subunit segmentation to cognitively healthy adults from the Open Access Series of Imaging Studies (OASIS-1)^21^ and examined associations of intracranial-volume-normalised subunit volumes with age and SES. We expected age to be negatively associated with subunit volumes, potentially in a region-specific manner, and we explored whether SES would show associations with subunits implicated in stress and neuroendocrine regulation.

## 2. Materials and Methods

### 2.1. Participants

Data were obtained from the OASIS-1 cross-sectional release, which comprises T1-weighted magnetic resonance imaging (MRI) scans and clinical data from 416 right-handed participants aged 18–96 years.^21^ Dementia status was assessed with the Clinical Dementia Rating (CDR)^22^ and global cognition with the Mini-Mental State Examination (MMSE).^23^ SES was rated with the Hollingshead Index of Social Position (1 = highest, 5 = lowest).^24^ To focus on healthy ageing, we excluded participants with CDR > 0 (n = 100) and participants without a CDR assessment (n = 181; aged 18–58 years), for whom healthy status could not be ascertained with the same criterion. Of the remaining 135 participants, two lacked segmentation output, yielding an analytic sample of 133 (SES available for 131; Figure 1A). Sample characteristics are presented in Table 1.

**Table 1.**
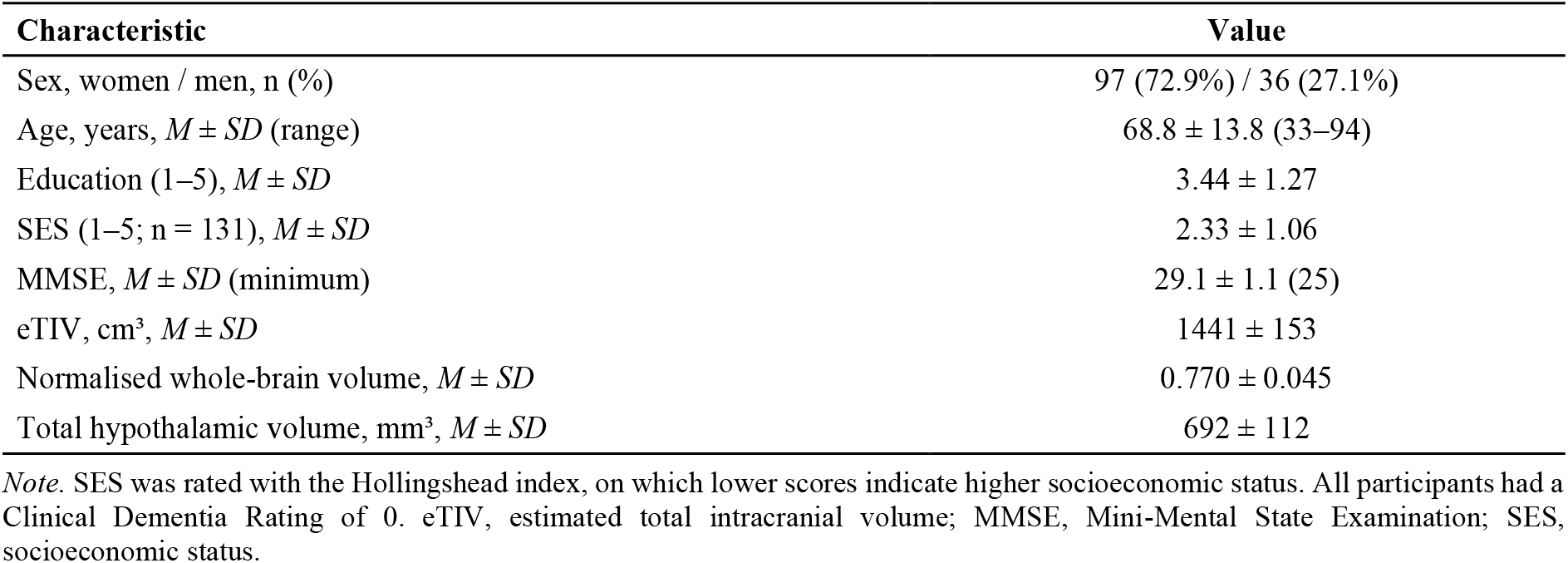
Demographic and clinical characteristics of the analytic sample (n = 133)

**Figure 1.**
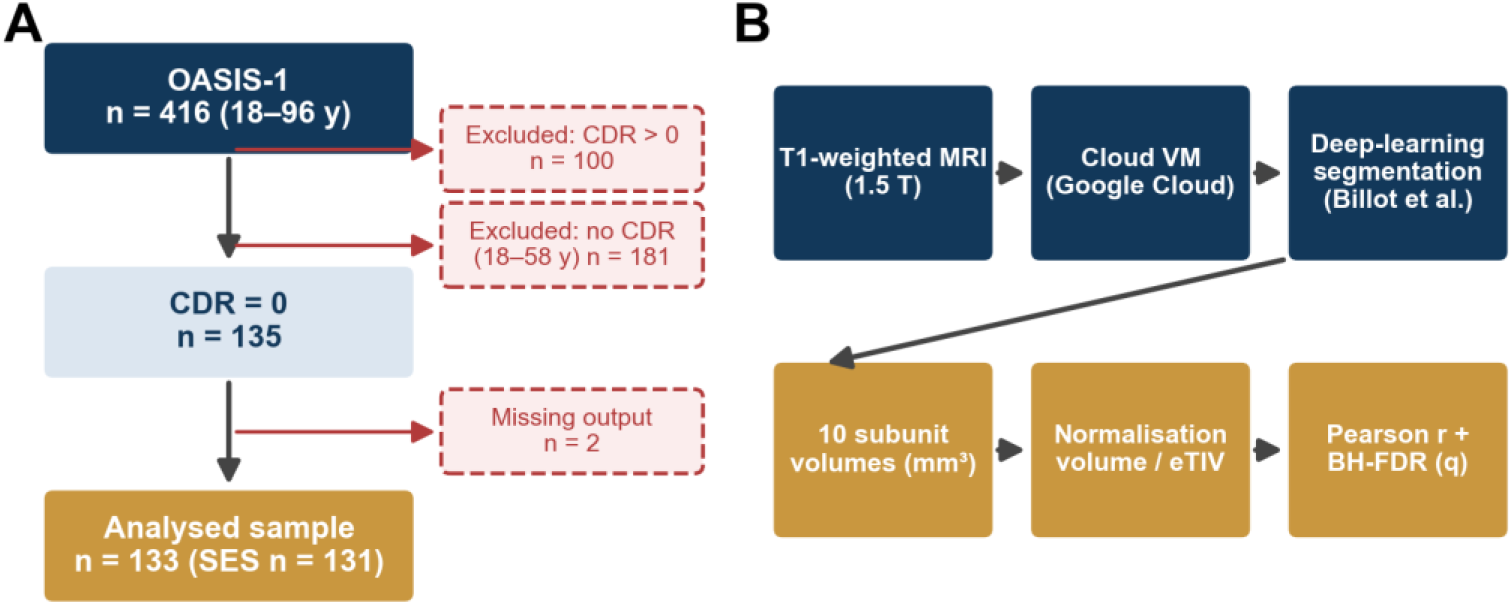
Study design. (A) Participant flow from the OASIS-1 cross-sectional dataset to the analytic sample. (B) Image-processing and analysis pipeline. CDR, Clinical Dementia Rating; eTIV, estimated total intracranial volume; BH-FDR, Benjamini–Hochberg false discovery rate.

### 2.2. MRI acquisition and hypothalamic segmentation

OASIS-1 images were acquired on a 1.5-T Siemens Vision scanner using an MP-RAGE sequence; for each participant, three to four individual scans were acquired and averaged to increase signal-to-noise ratio.^21^ Estimated total intracranial volume (eTIV) was provided with the dataset and was derived using atlas-based head-size normalisation.^25^ Hypothalamic segmentation was performed on the FreeSurfer-processed OASIS-1 data distributed by the OASIS project (11 archives), using the deep-learning hypothalamic subunit module (mri_segment_hypothalamic_subunits)^6^ of FreeSurfer 8.2.0, the latest release at the time of analysis; processing was executed on a Google Cloud virtual machine to ensure a uniform computational environment for all scans (Figure 1B). The module returns the volumes (mm^3^) of five subunits per hemisphere (Figure 2B) together with whole-hemisphere hypothalamic volumes. For visualisation, the hypothalamus was displayed on the MNI152 template using the CIT168 subcortical atlas (Figure 2A).^26^

**Figure 2.**
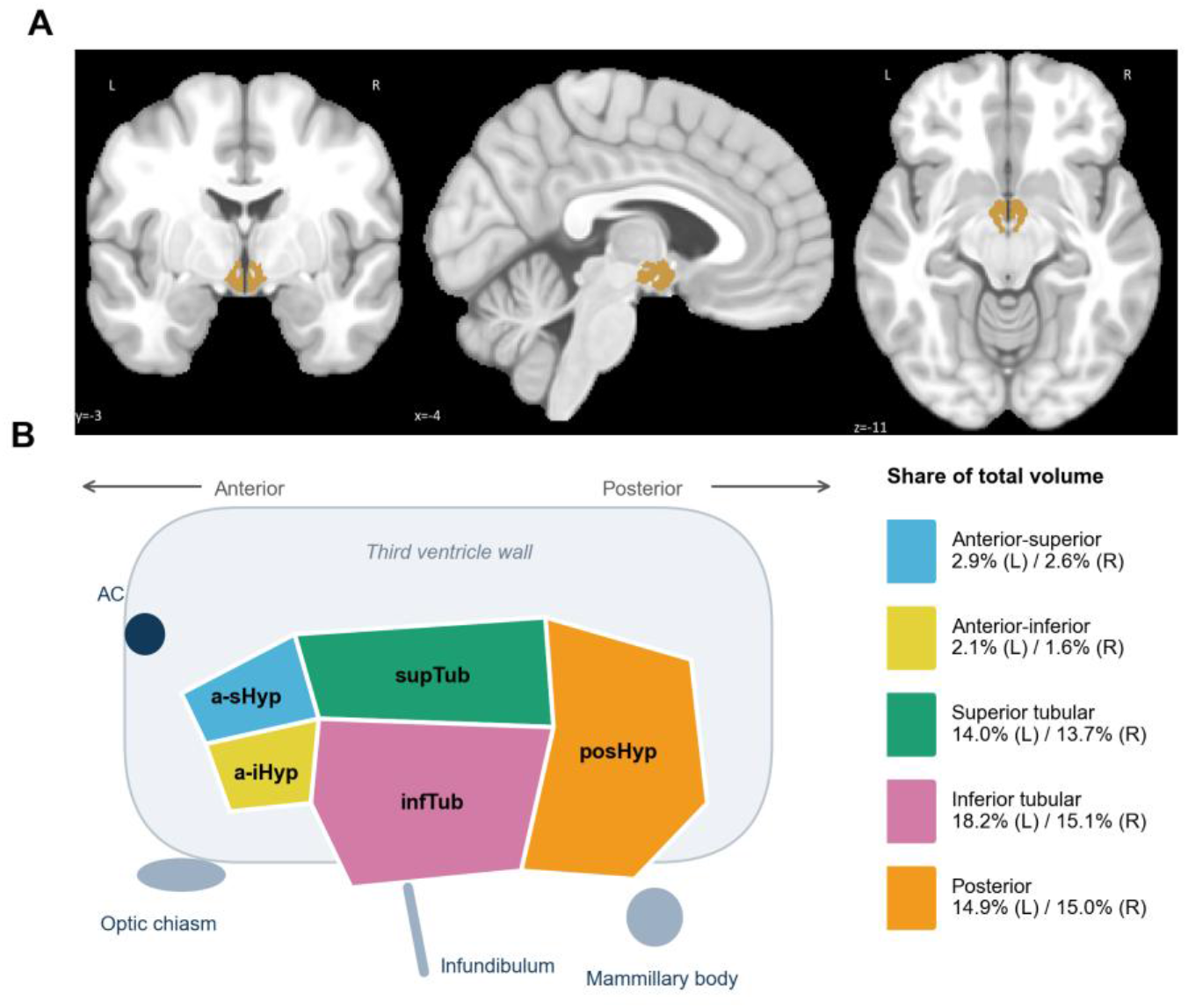
Anatomy of the hypothalamus and its subunits. (A) Location of the hypothalamus (gold) on coronal, sagittal and axial sections of the MNI152 template (CIT168 atlas). (B) Schematic sagittal representation of the five subunits segmented per hemisphere by the deep-learning model, with neighbouring landmarks. The legend gives the mean share of each subunit in total (bilateral) hypothalamic volume in the present sample. AC, anterior commissure; a-sHyp, anterior-superior; a-iHyp, anterior-inferior; supTub, superior tubular; infTub, inferior tubular; posHyp, posterior.

### 2.3. Statistical analysis

Each subunit volume was divided by eTIV to account for head size (reported as ×10^−3^). Associations of normalised volumes with age and SES were tested with Pearson correlations. For each predictor, the ten subunit tests (five subunits × two hemispheres) formed the primary family for Benjamini–Hochberg false discovery rate (FDR) correction,^27^ and corrected values are reported as *q*; whole-hemisphere and total volumes were examined as secondary outcomes. As sensitivity analyses, we computed partial correlations adjusting for sex (age analyses) and for age and sex (SES analyses), again with FDR correction. Post hoc power for the observed correlations was estimated with the Fisher *z* approximation (two-tailed), both at α = .05 and at α = .005, the Bonferroni-equivalent threshold for the ten subunit tests. Analyses were conducted in Python (pandas, SciPy, statsmodels); statistical significance was set at *q* < .05.

## 3. Results

### 3.1. Sample characteristics and subunit volumes

Mean total (bilateral) hypothalamic volume was 692 mm^3^, corresponding to approximately 0.05% of eTIV. The tubular and posterior subunits accounted for about 91% of hypothalamic volume, whereas the two anterior subunits together accounted for about 9% (Figure 2B; Table 2). Age and SES were weakly correlated (*r* = 0.14).

**Table 2.** Hypothalamic subunit volumes and their associations with age and socioeconomic status.

| **Subunit** | **Volume, mm³** | **%<br>total** | **Age *r*** | ***q*** | **Adj. *r*** | ***q*** | **SES *r*** | ***q*** | **Adj. *r*** | ***q*** |
|---|---|---|---|---|---|---|---|---|---|---|
| *Left hemisphere* |  |  |  |  |  |  |  |  |  |  |
| Anterior-superior | 19.1 ± 3.6 | 2.9 | −.135 | .204 | −.171 | .091 | .091 | .497 | .121 | .307 |
| Anterior-inferior | 14.0 ± 4.3 | 2.1 | .098 | .376 | .087 | .459 | .242 | .053 | .236 | .067 |
| Superior tubular | 95.1 ± 15.4 | 14.0 | −.080 | .449 | −.065 | .573 | .099 | .497 | .109 | .307 |
| Inferior tubular | 126.5 ± 23.4 | 18.2 | .038 | .734 | .044 | .686 | .101 | .497 | .096 | .345 |
| Posterior | 103.1 ± 20.2 | 14.9 | −.274 | .007 | −.290 | .004 | .063 | .531 | .111 | .307 |
| *Right hemisphere* |  |  |  |  |  |  |  |  |  |  |
| Anterior-superior | 17.8 ± 5.1 | 2.6 | −.173 | .092 | −.208 | .055 | .087 | .497 | .123 | .307 |
| Anterior-inferior | 11.5 ± 4.9 | 1.6 | .175 | .092 | .167 | .091 | .157 | .363 | .139 | .307 |
| Superior tubular | 95.3 ± 17.4 | 13.7 | −.179 | .092 | −.170 | .091 | .027 | .759 | .052 | .554 |
| Inferior tubular | 106.3 ± 26.8 | 15.1 | .016 | .859 | .026 | .770 | .076 | .497 | .072 | .459 |
| Posterior | 103.4 ± 21.0 | 15.0 | −.456 | < .001 | −.472 | < .001 | .074 | .497 | .161 | .307 |

*Note.* Volumes are *M* ± *SD* of raw volumes; correlations use eTIV-normalised volumes (age: n = 133; SES: n = 131). *q*, Benjamini–Hochberg FDR-corrected *p* value across the ten subunits for each predictor. Adjusted (Adj.) *r* values are partial correlations controlling for sex (age) or for age and sex (SES). Shaded rows highlight the posterior subunits.

### 3.2. Associations with age

Older age was associated with lower normalised volume of the right posterior subunit (*r* = −.456, *p* < .001, *q* < .001) and of the left posterior subunit (*r* = −.274, *p* = .001, *q* = .007; Figures 3 and 4A–B). Nominal associations were observed for the right anterior-superior (*r* = −.173), right superior tubular (*r* = −.179) and right anterior-inferior (*r* = .175) subunits, none of which survived correction (all *q* = .092). The posterior associations were robust to adjustment for sex (right: partial *r* = −.472, *q* < .001; left: partial *r* = −.290, *q* = .004). Among secondary outcomes, the right whole-hemisphere volume showed a nominal negative association with age (*r* = −.202, *p* = .020), whereas total hypothalamic volume did not (*r* = −.168, *p* = .053), indicating that age-related differences were more evident at the subunit level than in the hypothalamus as a whole.

**Figure 3.**
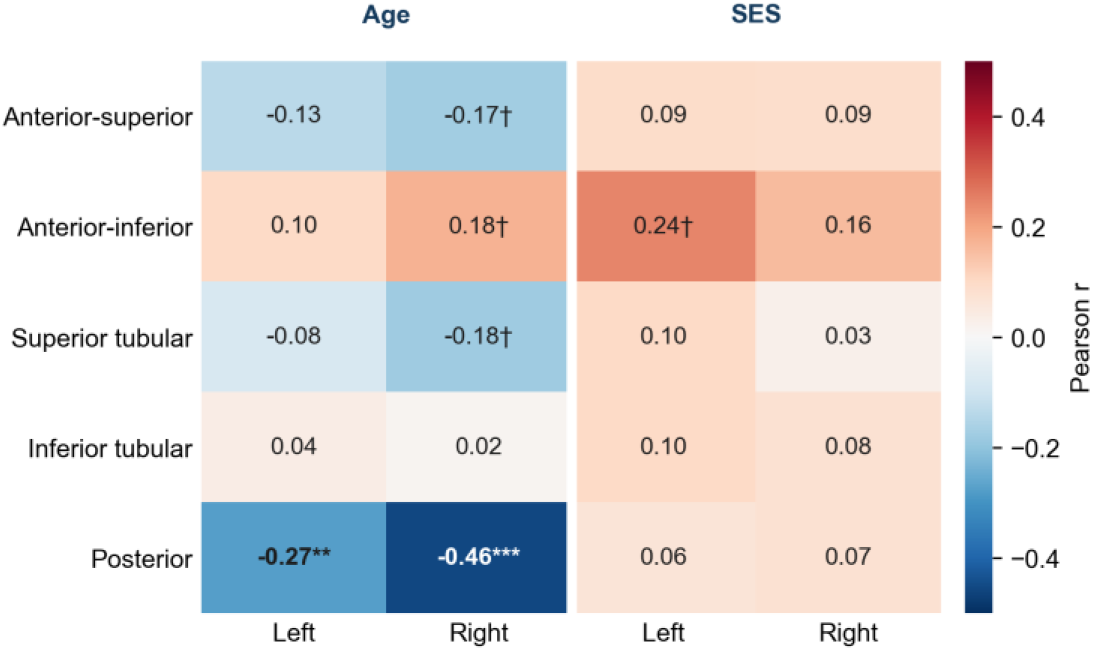
Correlation heat map of eTIV-normalised hypothalamic subunit volumes with age and SES. Cells show Pearson *r*; blue indicates negative and red positive associations. Asterisks denote Benjamini–Hochberg FDR-corrected significance within each predictor (*** *q* < .001, ** *q* < .01, * *q* < .05; † *q* < .10). Because lower Hollingshead scores denote higher status, positive SES correlations indicate larger volumes at lower socioeconomic status.

**Figure 4.**
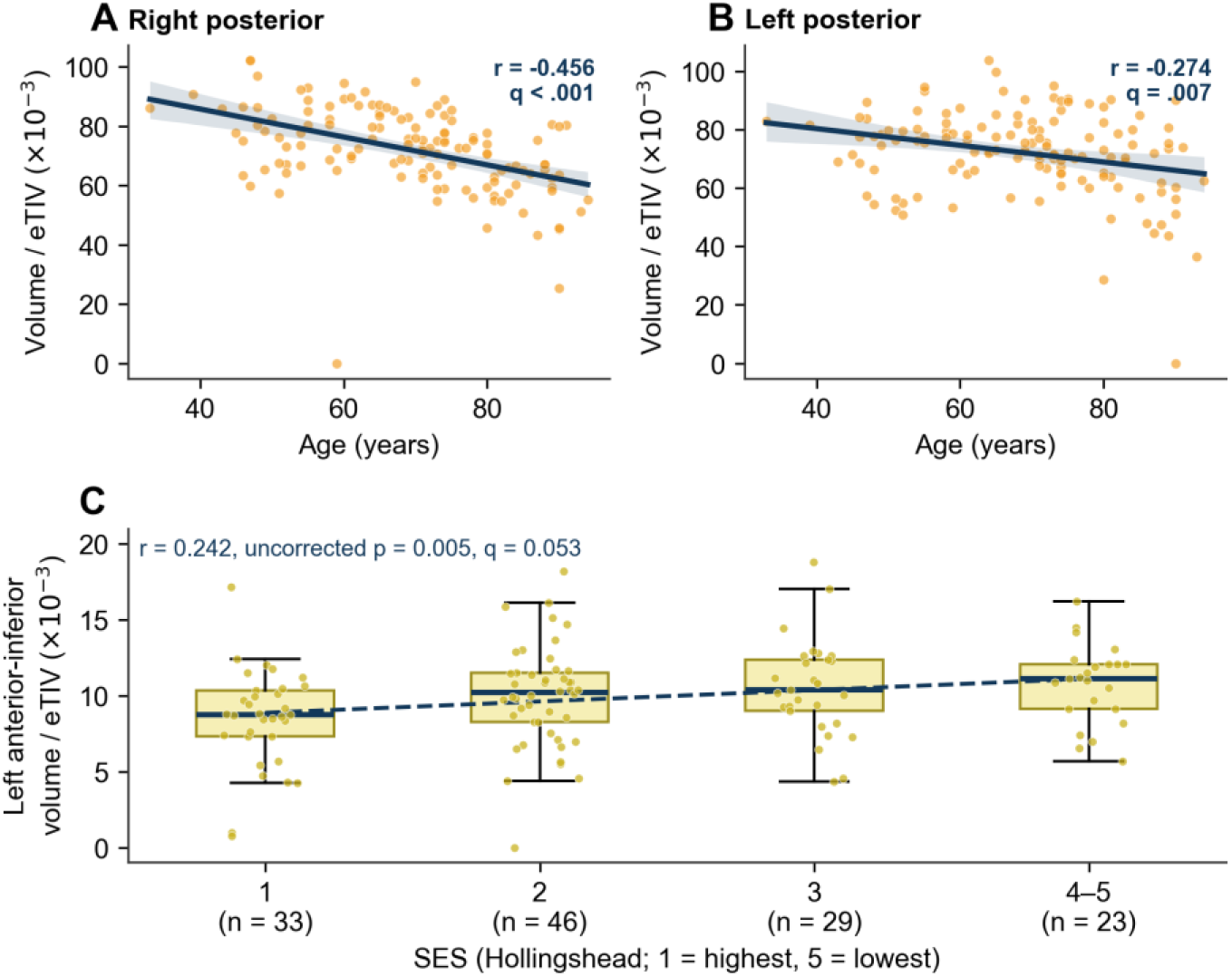
Age and SES associations with selected subunit volumes. (A, B) Normalised right and left posterior subunit volumes plotted against age; lines show linear fits with 95% bootstrap confidence bands. (C) Left anterior-inferior subunit volume by SES category (Hollingshead categories 4 and 5 combined for display); the dashed line shows the linear trend across all five categories. Values are volume/eTIV × 10^−3^.

### 3.3. Associations with socioeconomic status

SES showed a positive correlation with left anterior-inferior volume (*r* = .242, uncorrected *p* = .005), indicating larger volume at lower socioeconomic status; however, this association did not survive FDR correction (*q* = .053) and was similar after adjustment for age and sex (partial *r* = .236, *q* = .067; Figure 4C). No other subunit, whole-hemisphere or total volume was associated with SES (all *p* > .07). Post hoc power analysis showed that, with n = 131, power to detect *r* = .24 was .80 at α = .05 but only .50 at the multiplicity-adjusted threshold (α = .005); approximately 221 participants would be required for 80% power under correction. By contrast, power for the right posterior age association exceeded .99 even at α = .005.

## 4. Discussion

Using automated deep-learning segmentation in cognitively healthy adults, we found that age-related differences in hypothalamic volume were concentrated in the posterior subunits, most clearly in the right hemisphere, whereas total hypothalamic volume was not significantly associated with age. Lower SES showed only a statistical trend towards larger left anterior-inferior volume. These findings illustrate that subunit-level analysis can reveal regional effects that are diluted when the hypothalamus is treated as a single structure.

The posterior predominance of the age association converges with recent large-scale evidence. In two population-based cohorts comprising more than 50,000 participants, Xu and colleagues reported smaller anterior and posterior hypothalamic and mammillary body volumes, but larger tuberal volumes, with increasing age.^14^ Our results replicate the posterior finding with a different segmentation method and an independent dataset, and our nominal positive association between age and right anterior-inferior volume is directionally compatible with the heterogeneous, region-specific pattern described in that work. Functionally, the posterior subunit contains the mammillary bodies and portions of the lateral and tuberomammillary hypothalamus, which contribute to the ascending arousal system and to memory circuitry.^1,6,16^ Ageing is accompanied by fragmented sleep and weakened circadian rhythms, and fewer suprachiasmatic neurons have been linked to weaker rest–activity rhythms in older humans.^17^ It is therefore plausible that posterior hypothalamic volume differences reflect, or contribute to, age-related changes in arousal regulation; however, because no sleep measures were available, this interpretation remains a hypothesis for future work.

The SES trend should be interpreted cautiously, as it did not survive correction for multiple comparisons and the study was underpowered for effects of this size. Nevertheless, its direction is noteworthy. Socioeconomic disadvantage is associated with chronic stress, inflammation and differences in brain structure across the lifespan,^18–20^ and the anterior hypothalamus participates in neuroendocrine and circadian control.^1,2^ Larger hypothalamic volumes have also been reported in association with higher body mass index, possibly reflecting inflammatory or gliotic processes,^15^ and with childhood conduct problems, again involving the left anterior-inferior subunit.^12^ Because body mass index and early-life adversity were not available in OASIS-1, these factors may contribute to the observed trend and should be modelled in larger, better-characterised samples.

Methodologically, this study demonstrates that hypothalamic subunit volumetry can be performed reproducibly at scale using freely available tools in a cloud environment. As individual-difference effects in neuroimaging are typically modest, adequately powered and pre-registered studies are essential;^28,29^ automated pipelines such as the one used here make such sample sizes attainable for small subcortical structures.

### 4.1. Limitations

Several limitations should be acknowledged. First, the cross-sectional design precludes conclusions about causality or within-person volumetric change; age associations may partly reflect cohort effects. Second, the sample was modest, predominantly female and older, and was imaged at 1.5 T, which limits precision for very small structures; the anterior subunits, which together account for less than 10% of hypothalamic volume, are particularly susceptible to measurement error, and a small number of near-zero subunit estimates suggests occasional segmentation failure that would benefit from systematic visual quality control. Third, the primary analyses were bivariate; although sex-adjusted and age- and sex-adjusted sensitivity analyses yielded similar results, other potential confounders (e.g., body mass index, vascular risk factors, sleep) could not be assessed. Fourth, SES was indexed by a single ordinal measure. Finally, eTIV normalisation by proportion is one of several head-size correction approaches, and results may vary with alternative intracranial volume estimates.^30,31^

## 5. Conclusions

Automated deep-learning segmentation of the hypothalamus revealed that, among cognitively healthy adults, older age is associated with lower normalised volume of the posterior hypothalamic subunits, particularly on the right, whereas the association of lower socioeconomic status with larger left anterior-inferior volume remains a trend requiring confirmation. Longitudinal studies with larger samples, higher-field imaging and measures of sleep, metabolism and life adversity are needed to clarify the mechanisms and functional consequences of these regional differences.

## Declarations

### Ethics statement

This study is a secondary analysis of anonymised, publicly available data. OASIS-1 data were collected with institutional ethical approval and informed consent at Washington University in St. Louis; no additional data were collected for the present study.

## Data availability

OASIS-1 data are openly available at https://sites.wustl.edu/oasisbrains/. Analysis code is available from the corresponding author upon reasonable request.

## Acknowledgements

Data were provided by OASIS-1: Cross-Sectional (Principal Investigators: D. Marcus, R. Buckner, J. Csernansky, J. Morris; P50 AG05681, P01 AG03991, P01 AG026276, R01 AG021910, P20

MH071616, U24 RR021382). Preliminary findings were presented at the 9th BİLSEL International Harput Scientific Research Congress (17–18 October 2026, Elazığ, Türkiye).

## Author contributions (CRediT)

Barış Akar: Conceptualization, Methodology, Software, Formal analysis,

Visualization, Writing – original draft. Ümit Atasever: Supervision, Validation, Writing – review & editing.

## Use of generative AI

A generative AI tool was used to assist with drafting the text and producing figures from the authors’ analysis outputs. The authors reviewed and verified all content and take full responsibility for the manuscript.

